# *bla*_OXA-48_-like genome architecture among carbapenemase-producing *Escherichia coli* and *Klebsiella pneumoniae* in the Netherlands

**DOI:** 10.1101/2020.12.18.423568

**Authors:** Antoni P.A. Hendrickx, Fabian Landman, Angela de Haan, Dyogo Borst, Sandra Witteveen, Marga van Santen-Verheuvel, Leo M. Schouls, the Dutch CPE surveillance Study Group

**Author notes:** To whom correspondence should be send. contributed equally to this study (Antoni P.A. Hendrickx, Fabian Landman). list of participants can be found at the end of this article.

## Abstract

Carbapenem-hydrolyzing enzymes belonging to the OXA-48-like group are encoded by *bla*_OXA-48_-like alleles and are abundant among *Enterobacterales* in the Netherlands. Therefore, the objective was to investigate the characteristics, gene content, and diversity of the *bla*_OXA-48_-like carrying plasmids and chromosomes of *Escherichia coli* and *Klebsiella pneumoniae* collected in the Dutch national surveillance from 2014-2019 in comparison with genome sequences retrieved from 29 countries. By combining short-read and long-read sequencing, 47 and 132 complete *bla*_OXA-48_-like plasmids were reconstructed for *E. coli* and *K. pneumoniae*, respectively. Distinct plasmid groups designated as pOXA-48, pOXA-181, and pOXA-232 were identified in the Netherlands and varied in size, % G+C, presence of antibiotic resistance genes, replicons and gene content. The pOXA-48 plasmids were variable, while pOXA-181 and pOXA-232 plasmids were conserved. A group of non-related pOXA-48 plasmids contained different resistance genes, non-IncL type replicons or carried no replicons. *K. pneumoniae* isolates carrying *bla*_OXA-48_ or *bla*_OXA-232_ were mostly resistant, while *E. coli bla*_OXA-48_, *bla*_OXA-181_ and chromosomal *bla*_OXA-48_ or *bla*_OXA-244_ isolates were mostly sensitive for meropenem. Analysis of chromosomally localized *bla*_OXA-48_-like alleles revealed that these were flanked by a direct repeat (DR) upstream of IS1R, which were found at multiple locations in the chromosome of distinct genetic backgrounds. In conclusion, the overall *bla*_OXA-48_-like plasmid population in the Netherlands is conserved and similar to that reported for other countries, although a highly diverse *bla*_OXA-48_-like plasmid subgroup was present. Chromosomally encoded *bla*_OXA-48_-like alleles are from distinct genetic backgrounds and occurs at variable positions containing the DR, thereby indicating multiple independent transpositions.

**Importance:** OXA-48-type of carbapenem hydrolyzing enzymes encoded by *bla*OXA-48-like genes from transmissible plasmids or chromosomes of *Escherichia coli* and *Klebsiella pneumoniae* have spread world-wide and are of concern. Dissecting the *bla*OXA-48-like genome architecture at the molecular level by combining short-read and long-read sequencing will lead to understanding trends in the plasmid reservoir of *E. coli* and *K. pneumoniae* in the Netherlands and may enhance future international pathogen surveillance.

## Introduction

Antimicrobial resistance (AMR) has dispersed among the family of *Enterobacterales* and is a major concern for both hospitalized and non-hospitalized patients (1). In carbapenemase-producing *Enterobacterales* (CPE), genes encoding carbapenemases are often located on transmissible plasmids that shuttle between bacterial strains of the same species, but also between distinct bacterial species and often confer resistance to carbapenem antibiotics (2, 3). The predominant CPE species in the Netherlands from 2014 to 2019 were *Klebsiella pneumoniae* (43%), *Escherichia coli* (30%) and *Enterobacter cloacae* complex (13%) (4). Carbapenemases are classified in Ambler classes A (*i*.*e*., KPC-types), B (*i*.*e*., IMP-, NDM- and VIM-types), and D (OXA β-lactamases) type of carbapenem antibiotic degrading enzymes (5). The KPC, NDM, IMP, VIM and certain OXA-like enzymes are the most commonly identified variant carbapenemases that have spread world-wide among *Enterobacterales*, including *E. coli* and *K. pneumoniae* (6). The *bla*_OXA-48_-like genes make up the most prevalent carbapenemase-encoding genes found in *Enterobacterales* in the Netherlands (44%), followed by *bla*_NDM_ (27%) (4). The OXA-48-like carbapenemases are encoded by the *bla*_OXA-48_, *bla*_OXA-162_, *bla*_OXA-181_, *bla*_OXA-204_, *bla*_OXA-232_, and *bla*_OXA-244_ genes. Other OXA-48-likes, such as OXA-245, OXA-484, and OXA-519 are less often reported groups of carbapenemases (6). The distinction between the OXA-48-like carbapenemases is based on one to five specific amino acid substitutions in the β5-β6-loop of the enzyme that can impact the efficiency of carbapenem hydrolysis (6–8). OXA-181 differs from OXA-48 by four amino acids substitutions (Thr104Ala, Asn110Asp, Glu168Gln and Ser171Ala) yet have both comparable carbapenem hydrolytic activity (9). OXA-232 differs from OXA-48 by five amino acid substitutions, four are identical to the differential OXA-181 mutations, however OXA-232 contains an additional Arg214Ser substitution (10). OXA-244 differs only by a single Arg214Gly mutation from OXA-48 and the OXA-244 together with OXA-181 enzymes have reduced carbapenem hydrolyzing activity (11).

The most common plasmids that harbor *bla*_OXA-48_ belong to the IncL/M family, which are conjugative and have been described for *E. coli* and *K. pneumoniae* (12–15). The *bla*_OXA-181_ gene is located on plasmids containing the *qnrS1* gene encoding for quinolone resistance and either ColE2, IncX3, IncN1 or IncT type of replicons (16, 17). Plasmids containing *bla*_OXA-232_ have the ColE-type of replicon and the backbone is identical to *bla*_OXA-181_ containing plasmids (10). The *bla*_OXA-244_ gene is located on an IncL plasmid and is suggested to be originated from *bla*_OXA-48_ by a point mutation, which possibly occurred during integration in the *E. coli* ST38 chromosome (6, 11, 15). Chromosome encoded OXA-48-like carbapenemases have been described previously in globally disseminated *E. coli* and *K. pneumoniae* (15, 18, 19). In these chromosomes, the *bla*_OXA-48_-like gene has been found to be inserted at various chromosomal locations (18).

The global emergence of the carbapenem-hydrolyzing OXA-48 enzyme and OXA-48-like descendants on transmissible plasmids warrants a national surveillance. Currently, a paradigm shift occurs in national reference laboratories from next-generation sequencing (NGS) towards third generation long-read sequencing (TGS). This allows an in-depth study of CPE antibiotic resistance-plasmid biology and plasmid transmission within and between health care institutions and countries, respectively. Therefore, the major goal of this study was to investigate the characteristics and contents of *E. coli* and *K. pneumoniae* plasmids and chromosomes carrying *bla*_OXA-48_-like genes obtained from isolates submitted to the Dutch national CPE surveillance program in a global context using a combination of NGS and TGS.

## Results

### Resistance to meropemen of CPE carrying *bla*_OXA*-*48_-like plasmids

From 2014 till 2019, the National Institute for Public Health and the Environment (RIVM) received 1503 carbapenemase-producing *Enterobacterales*, of which the majority (*n*= 1106) were *E. coli* (*n*= 461) and *K. pneumoniae* (*n*= 645). PCR revealed that 272 *E. coli* and 338 *K. pneumoniae* isolates carried *bla*_OXA-48_-like alleles. Only the first submitted *E. coli* or *K. pneumoniae* isolate with a *bla*_OXA-48_-like allele per person in this study period was included. Therefore, 537 carbapenemase-producing *E. coli* (*n*= 230) and *K. pneumoniae* (*n*= 307) isolates were sequenced by NGS (Table S1, Suppl. file 1). The majority of the *E. coli* isolates were carrying *bla*_OXA-48_, *bla*_OXA-181_, and *bla*_OXA-244_ alleles and had MICs for meropenem that were below the clinical breakpoint of 2 mg/L for sensitivity according to EUCAST (206/230; 89.6%) (Table 1). Only 2/157 (1.3%) of the *E. coli* isolates with *bla*_OXA-48_ reached the clinical breakpoint for resistance (>8 mg/L) to meropenem. The *bla*_OXA-244_ allele was found predominantly in *E. coli* (30/32; 93.8%) and was associated with a low MIC for meropenem. *K. pneumoniae* carried mostly *bla*_OXA-48_, *bla*_OXA-181_, and *bla*_OXA-232_ alleles, of which the *bla*_OXA-48_ allele was associated with resistance to meropenem (63/307; 20.5%). The *bla*_OXA-181_ allele was found in both *E. coli* and *K. pneumoniae*, and conferred resistance to meropenem in 4/21 (19%) of the *K. pneumoniae* isolates and 1/36 (2,8%) of the *E. coli* isolates. The *bla*_OXA-232_ allele was exclusively found in *K. pneumoniae* and none of these isolates were meropenem sensitive (R, 17/19; 89.5%, I, 2/19; 10.5%). Combinations of *bla*_OXA-48_-like alleles with either *bla*_NDM-1_ or *bla*_NDM-5_ resulted in high MICs for meropenem. For all *bla*_OXA-48_-like alleles and double allele combinations *K. pneumoniae* was more resistant (123/307; 40.1%) than *E. coli* (9/230; 3.9%). Due to initial limited resources, a subset (220/537; 41%) of the isolates submitted in 2018 and 2019 were sequenced with Nanopore long-read sequencing enabling the reconstruction of 47 and 132 complete *bla*_OXA-48_-like plasmids for *E. coli* and *K. pneumoniae*, respectively (Table 2).

**Table 1.**
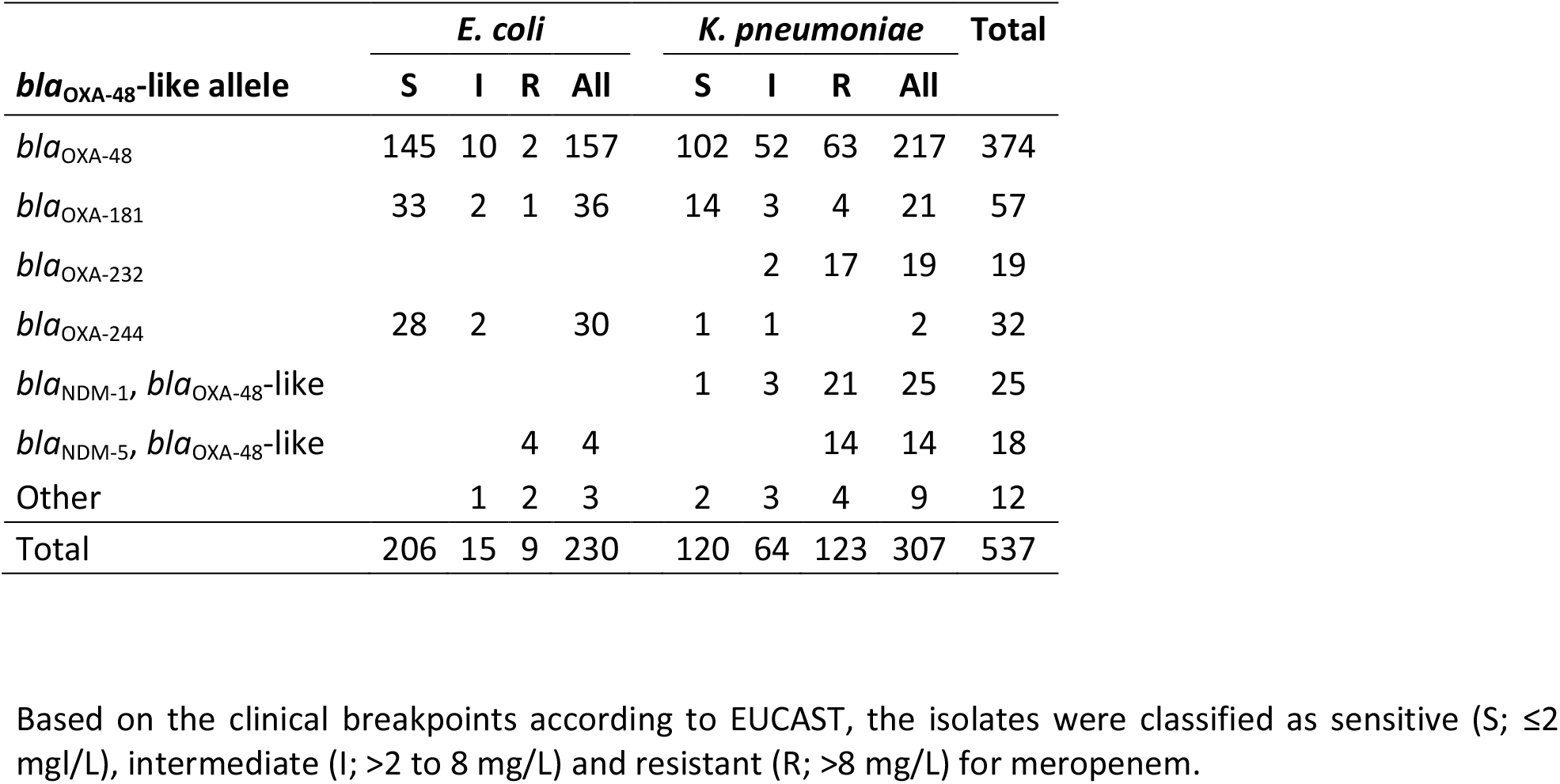
Resistance to meropenem per *E. coli* or *K. pneumoniae* isolate carrying *bla*_OXA-48_-like alleles 2014-2019.

| *bla*<sub>OXA-48</sub>-like allele | *E. coli* |  |  |  | *K. pneumoniae* |  |  |  | Total |
| --- | --- | --- | --- | --- | --- | --- | --- | --- | --- |
|  | S | I | R | All | S | I | R | All |  |
| *bla*<sub>OXA-48</sub> | 145 | 10 | 2 | 157 | 102 | 52 | 63 | 217 | 374 |
| *bla*<sub>OXA-181</sub> | 33 | 2 | 1 | 36 | 14 | 3 | 4 | 21 | 57 |
| *bla*<sub>OXA-232</sub> |  |  |  |  |  | 2 | 17 | 19 | 19 |
| *bla*<sub>OXA-244</sub> | 28 | 2 |  | 30 | 1 | 1 |  | 2 | 32 |
| *bla*<sub>NDM-1</sub>, *bla*<sub>OXA-48</sub>-like |  |  |  |  | 1 | 3 | 21 | 25 | 25 |
| *bla*<sub>NDM-5</sub>, *bla*<sub>OXA-48</sub>-like |  |  | 4 | 4 |  |  | 14 | 14 | 18 |
| Other |  | 1 | 2 | 3 | 2 | 3 | 4 | 9 | 12 |
| Total | 206 | 15 | 9 | 230 | 120 | 64 | 123 | 307 | 537 |

Based on the clinical breakpoints according to EUCAST, the isolates were classified as sensitive (S; ≤2 mg/L), intermediate (I; >2 to 8 mg/L) and resistant (R; >8 mg/L) for meropenem.

**Table 2.** *bla*_OXA-48_-like plasmids and chromosomes analyzed in this study.

| Plasmid/chromosome | Species | Carbapenemase allele |  |  |  | Total |
| --- | --- | --- | --- | --- | --- | --- |
|  |  | *bla*<sub>OXA-48</sub> | *bla*<sub>OXA-181</sub> | *bla*<sub>OXA-232</sub> | *bla*<sub>OXA-244</sub> |  |
| Plasmids | *E. coli* | 30 | 16 | 1 |  | 47 |
|  | *K. pneumoniae* | 108 | 10 | 14 |  | 132 |
| Plasmids NCBI | *E. coli* | 14 | 35 |  |  | 49 |
|  | *K. pneumoniae* | 81 | 10 | 22 | 1 | 114 |
| Chromosomes | *E. coli* | 30 |  |  | 10 | 40 |
|  | *K. pneumoniae* | 4 |  |  |  | 4 |
| Chromosomes NCBI | *E. coli* | 6 |  |  | 1 | 7 |
|  | *K. pneumoniae* | 1 |  |  |  | 1 |
| Total |  | 274 | 71 | 37 | 12 | 394 |

Plasmids included in this study are complete and circular only, while the chromosomes were either circular or linear DNA.
NCBI indicates plasmids or chromosomes retrieved from the National Center for Biotechnology Information.

### *bla*_OXA-48_-like plasmids cluster in distinct genogroups

Comparison of the *bla*_OXA-48_-like plasmid sequences retrieved from the Netherlands with internationally reported *bla*_OXA-48_-like plasmids revealed clustering of the plasmids in a pOXA-232 group, a pOXA-181 group and five distinct pOXA-48 groups (Fig. 1a). A number of plasmids did not cluster with any of the other plasmids and were designated as the “non-cluster” group. Plasmids identified in the Netherlands were similar to internationally reported plasmids that were obtained from 29 different countries from North and South America, Europe, Asia and Oceania (Suppl. File 1). In general, there was a paucity of antibiotic resistance genes in most of the *bla*_OXA-48_-like containing plasmids (Fig. 1b). Unweighted pair group method with arithmetic mean (UPGMA) clustering based on plasmid sequence comparison showed that the pOXA-232 plasmids containing the ColKP3 replicon were highly conserved (96-100% similarity). With 6.2 kb in size, the pOXA-232 plasmids were the smallest *bla*_OXA-48_-like plasmids and carried a single replicon, but had the highest average %G+C content of 52.2% (Fig. 2). In contrast, pOXA-181 plasmids carried the *qnrS1* allele and ColKP3 and IncX3 replicons and were also conserved (90-100%)(Fig. 1a + b). The pOXA-181 plasmids were on average 51.3 kb in size and have the lowest %G+C content of 46.4% (Fig. 2). Despite the high sequence conservation of pOXA-181 and pOXA-232 plasmids, they were found in CPE with distinct chromosomal backgrounds (Table S2). The largest and most variable group comprised *bla*_OXA-48_ containing plasmids with an IncL/M(pOXA-48) type of replicon and could be divided into five subgroups pOXA-48-1 to pOXA-48-5. The sequence conservation among pOXA-48-1 plasmids ranged from 80% to 100% (Fig. 1a). pOXA-48-1 plasmids were on average 64 kB with a %G+C content of 51.2% and differed only from pOXA-48-2 plasmids by 0.1 kb. pOXA-48-3 was characterized by the presence of the aminoglycoside resistance genes *aph(3’)-Ib, aph(3’)-VIb, aph(6’)-Id* and the ESBL gene *bla*_CTXM-14b_ (Fig. 1b, Fig. 2a). pOXA-48-3 plasmids resembled pOXA-48-5 plasmids, however most of the pOXA-48-5 plasmids lacked the *aph(6’)-Id* gene and contained a distinct IncL/M(pMU407) replicon (Fig. 1b). pOXA-48-4 plasmids lacked these aminoglycoside resistance genes and these plasmids were smaller in size. pOXA-48-3 and pOXA-48-5 had on average four AMR genes, one replicon per plasmid, a highly similar %G+C content of 50.9% and 50.7%, respectively, but differ 3.4 kB in size (Fig. 2a). Non-cluster *bla*_OXA-48_-like plasmids were distinct from those in the other groups and carried a wide variety of AMR genes resulting in distinct resistomes. These plasmids had either non-IncL/M type of replicons *e*.*g*. IncR, IncY, IncF, or IncA or no known replicons (Fig. 1a + b). In addition, they had plasmid sizes that differed from the those in the different plasmid groups and %G+C content and predominantly originated from isolates from the Netherlands (Fig. 2).

**Figure 1.**
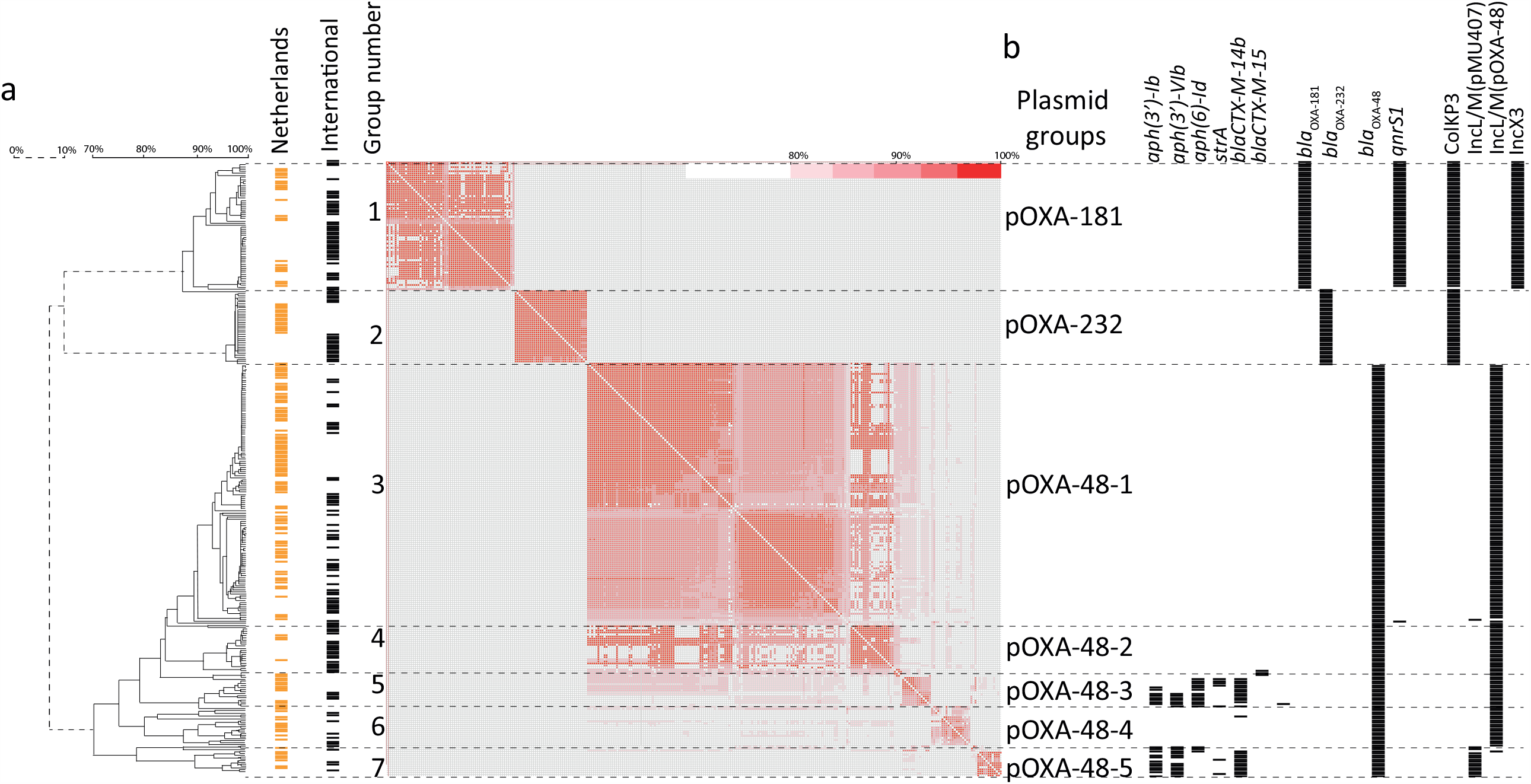
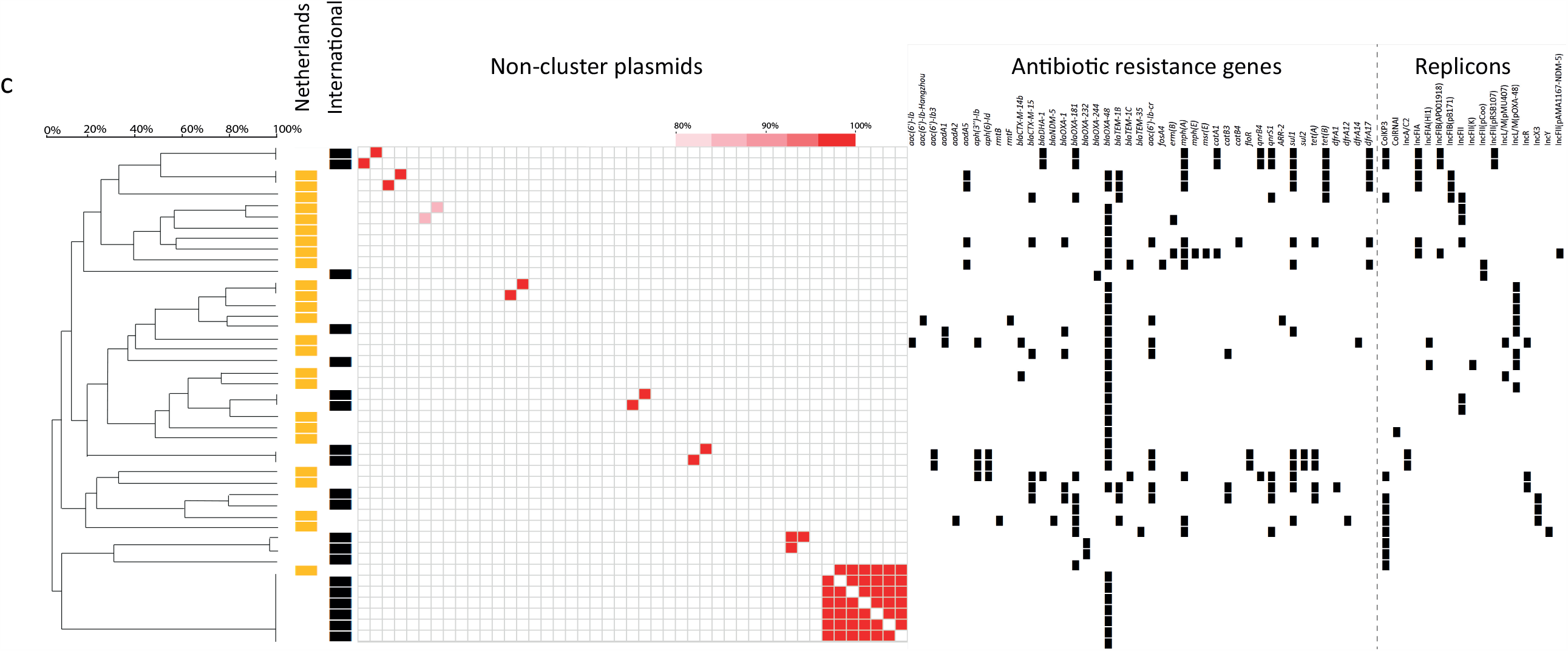
Genetic clustering of *bla*_OXA-48_-like plasmids. (a) UPGMA clustering based on plasmid DNA sequences revealed seven distinct groups of *bla*_OXA-48_-like plasmids. These groups were designated as pOXA-xxx, e.g. pOXA-48-1. Plasmids retrieved from the Netherlands are indicated in orange and international plasmids in black. Group numbers are indicated. A heatmap shows the percentage of sequence identity, where red is 100% identical and white 0% identical. (b) The presence of AMR genes and replicons among the plasmids is indicated with black squares. Plasmids are depicted in rows and the AMR genes and replicons in columns. (c) Similar as (a) and (b) for the non-cluster plasmids.

**Figure 2.**
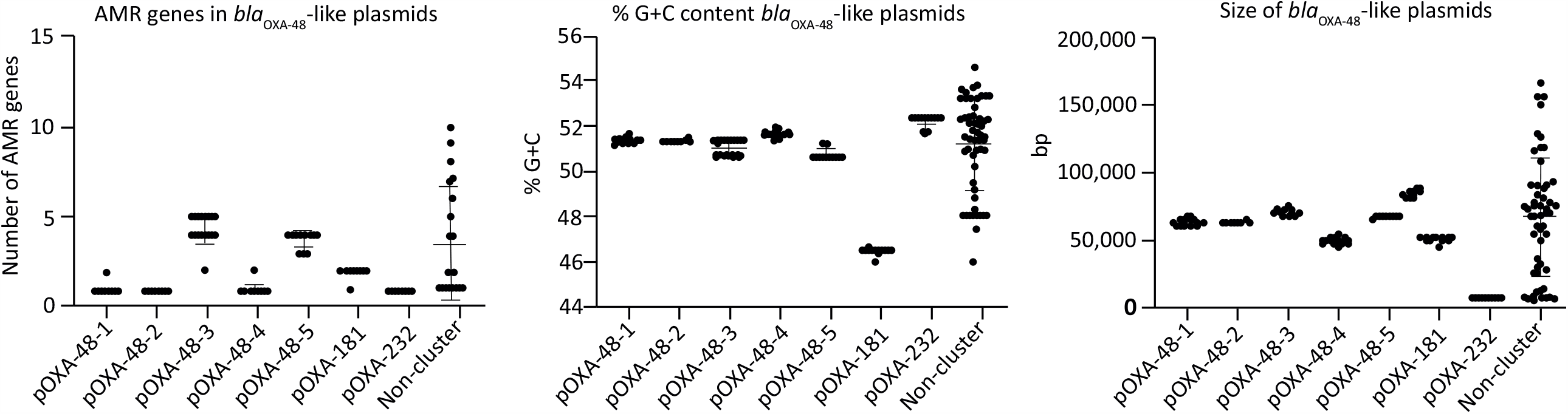
*bla*_OXA-48_-like plasmids have distinct characteristics. (a) Graphs show the distinct characteristics *e*.*g*. the number of AMR genes, the size in kilo base pairs and the %G+C content of the distinct pOXA-48-like groups of plasmids. Each dot represents a value per plasmid. Scale bars indicated the variation per group.

### Gene content determines distinct *bla*_OXA-48_-like plasmid architecture

Analysis of the gene content of representative plasmids from the seven distinct plasmid groups revealed a group-associated gene content (Fig. 3). pOXA-48 plasmids had conserved plasmid regions, designated as region 1, 2 and 3 and a central variable region (VR) which displayed variability in gene content and length (Fig. 3a). Plasmid region 2 was absent in pOXA-48-4 plasmids. The variations in pOXA-48 plasmid gene content such as the presence or absence of AMR genes shaped the primary *bla*_OXA-48_-like plasmid architecture, and varied among the different plasmid groups. While the pOXA-48-1, pOXA-48-2, pOXA-48-3 and pOXA-48-5 groups contained the *klcA* anti-restriction gene and a putative conjugation system, these features were absent in the pOXA-48-4, pOXA-181 and pOXA-232 plasmid groups, indicating these plasmids are non-conjugative (Fig. 3b). In pOXA-48-4 plasmids, the *tra* conjugation system is incomplete, while pOXA-48-5 plasmids contain a full conjugation system. pOXA-48 plasmids contained the PemI antitoxin, while the pOXA-181 and pOXA-232 plasmids did not. pOXA-181 plasmids carried a *virB2-virB3-virB9-virB10-virB11* type IV secretion system, while the other pOXA-48 plasmids lacked this system. IS1 family transposases IS1R, IS1D, and IS4 family transposase IS10A were predominantly found in the pOXA-48 plasmids, and pOXA-181 plasmids were characterized by a variety of Tn3 family transposases. pOXA-232 plasmids did not contain IS or Tn3 elements.

**Figure 3.**
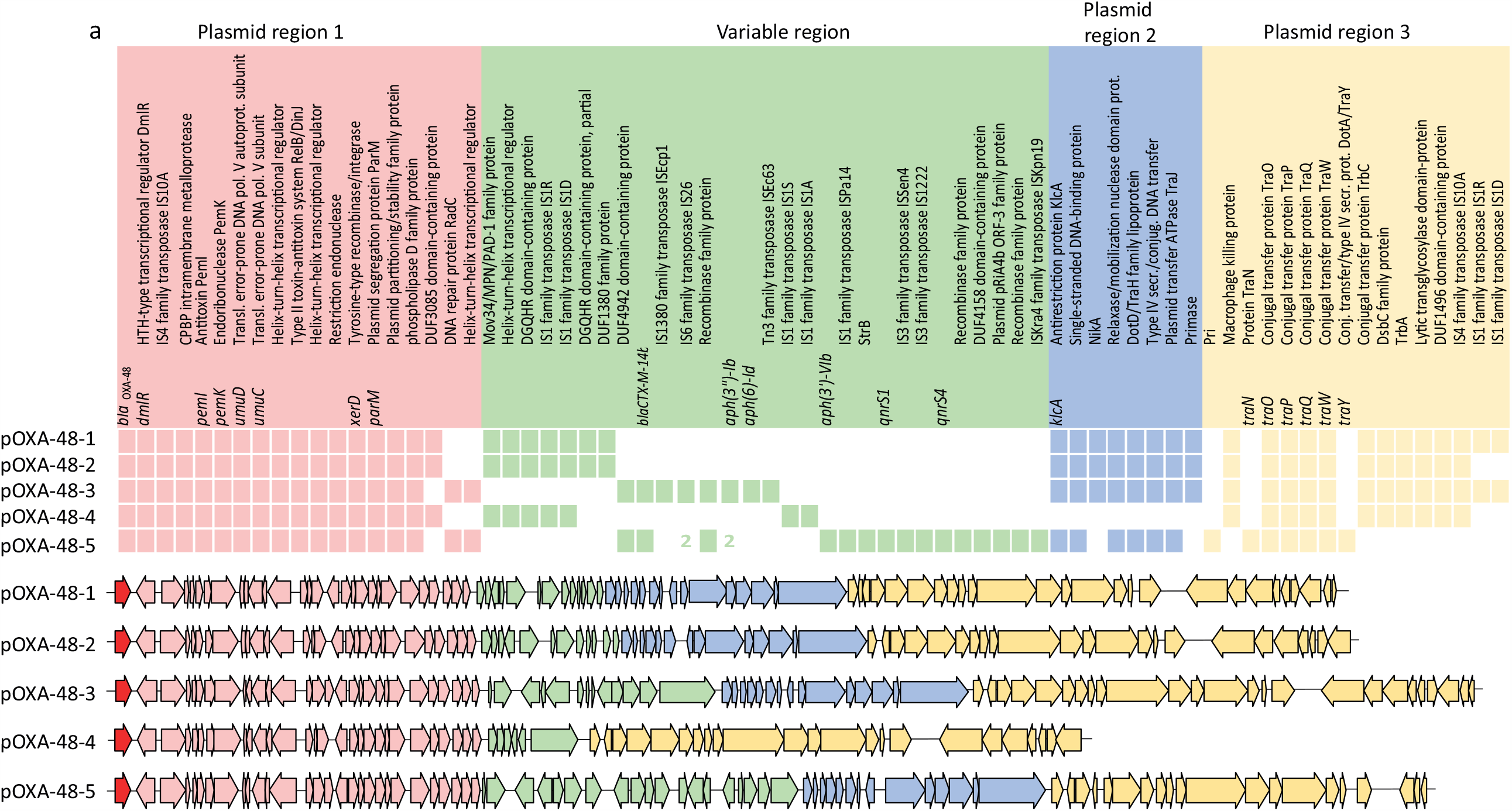
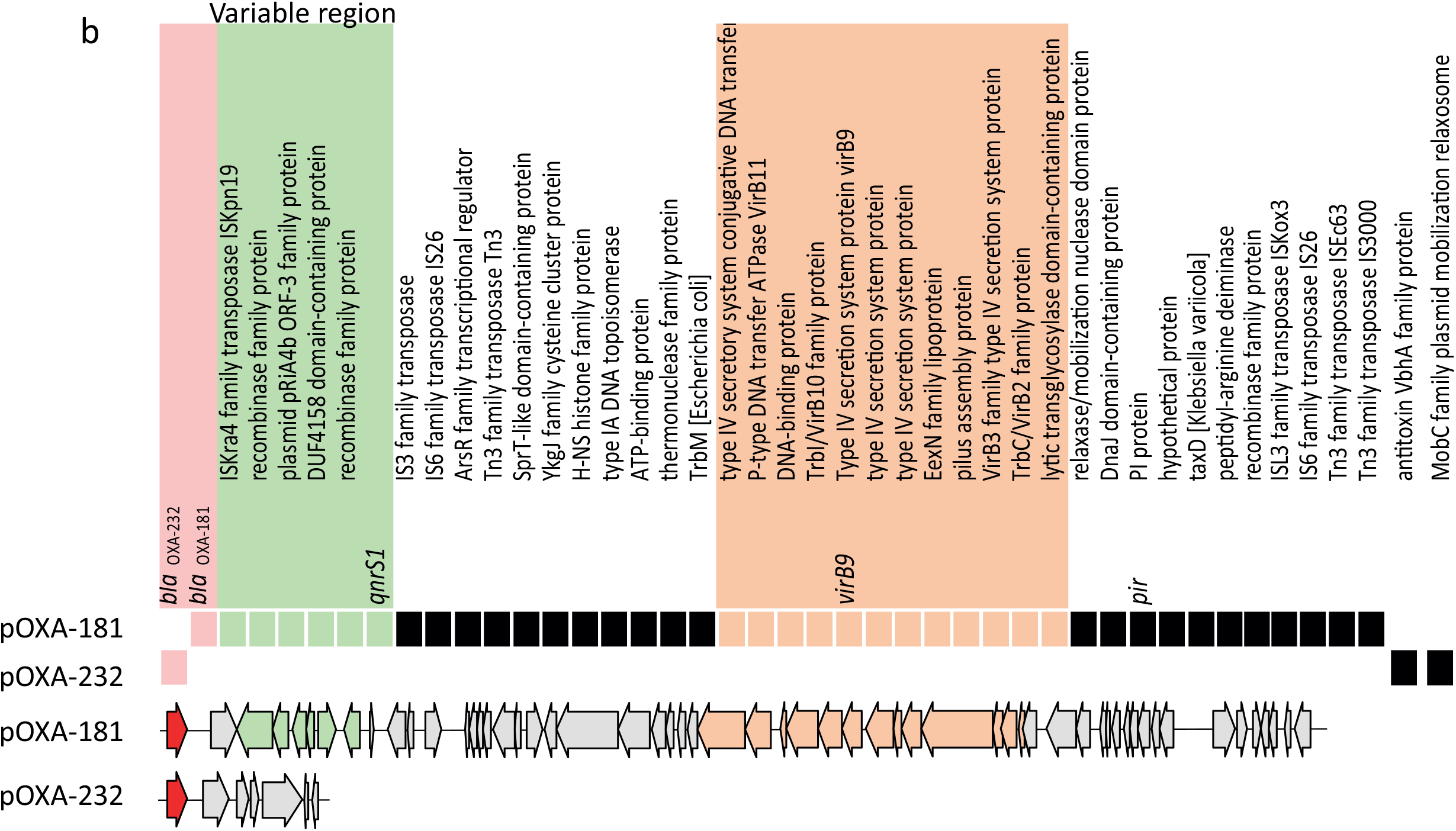
*bla*_OXA-48_-like plasmid architecture. (a) Diversity in pOXA-48-1 to pOXA-48-5 plasmid gene content. Complete plasmids were visualized in a linear way with the *bla*_OXA-48_-like allele at starting position 1. The presence and absence of genes is indicated among representative plasmids from the plasmid groups. Colors indicated different groups of genes corresponding to different regions in the plasmid, or the variable region. Plasmid regions are labelled above the plasmid sequence. (b) Similar as (a) but displays diversity in pOXA-181 and pOXA-232 plasmid gene content.

### Distribution of isolates harboring plasmid or chromosomally-localized *bla*_OXA-48_ and *bla*_OXA-244_ alleles

A fraction of the *bla*_OXA-48_ (30/230; 13%) and *bla*_OXA-244_ (10/230; 4.3%) alleles were located in the chromosomes of *E. coli* isolates, respectively (Table 2). Chromosomal *bla*_OXA-48_ or *bla*_OXA-244_ occurred in *E. coli* isolates with the MLST sequence type ST38, ST69, and ST127 among other STs (Fig. 4a, Table S2). The STs were all unrelated and were multiple locus variants from ST38. The chromosome-localized *bla*_OXA-48_ and *bla*_OXA-244_ were non-randomly distributed in the minimum spanning tree (MST) and restricted to specific STs (Fig. 4a, Table S2). In contrast, plasmid-localized *bla*_OXA-48_ occurred in *E. coli* isolates from a variety of non-related STs and were found randomly dispersed among the MST, except in ST38, ST69 and ST127. In four *K. pneumoniae* isolates *bla*_OXA-48_ was found to be integrated in the chromosome (4/307; 1.3%), while none of the *bla*_OXA-181_ and *bla*_OXA-232_ alleles were located chromosomally. *K. pneumoniae* with either chromosome- or plasmid-localized *bla*_OXA-48_-like were randomly distributed in the MST (Fig. 4b). The presence of the *bla*_OXA-48_ allele in the *E. coli* ST38 chromosomes was associated with the presence of the macrolide, trimethoprim and sulphonamide AMR genes *mph(A), dfrA*, and *sul*, while ST69 and ST127 were lacking the *dfrA* and *sul* genes (Fig. 4c). In contrast to *E. coli* ST38, the *bla*_OXA-48_ containing *K. pneumoniae* chromosomes were mostly devoid of AMR genes, with the exception of the fosfomycin and quinolone resistance genes *fosA, oqxA* and *oqxB*.

**Figure 4.**
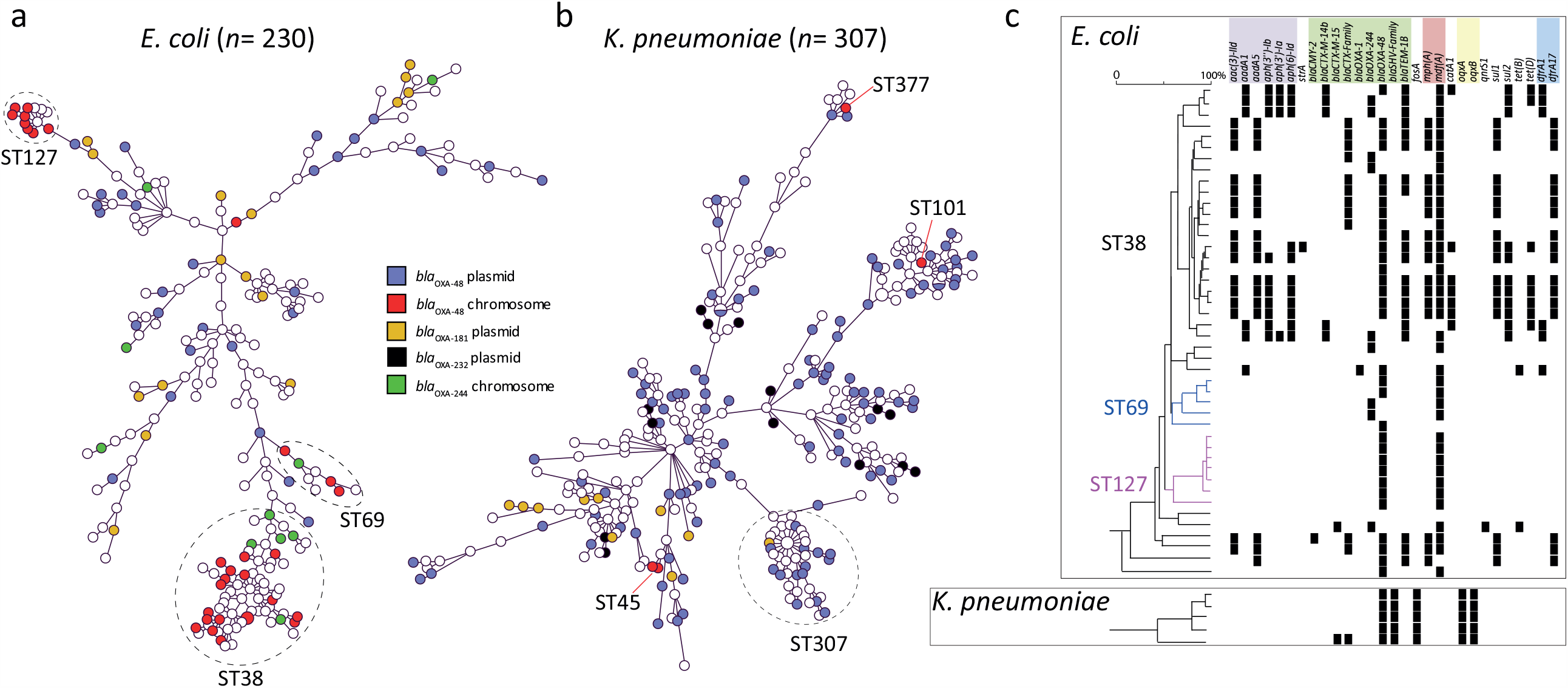
Distribution of chromosome- or plasmid-localized *bla*_OXA-48_ or *bla*_OXA-244_. (a) Minimum spanning tree of *E. coli* in which chromosome- or plasmid-localized *bla*_OXA-48_ or *bla*_OXA-244_ alleles are indicated by different colors. (b) Similar as in (a) but for *K. pneumoniae*. (c) The presence of AMR genes among the chromosomes analyzed in this study is indicated with black squares. Chromosomes are depicted in rows and the AMR genes in columns. Antibiotic classes are indicated above the AMR genes in different colors.

### Architecture of chromosome-localized *bla*_OXA-48_ or *bla*_OXA-244_ allelic regions

In *E. coli*, the *bla*_OXA-48_ or *bla*_OXA-244_ alleles were positioned in distinct regions in the chromosome relative to *dnaA* (Fig. 5a, Table 3). Chromosomally residing *bla*_OXA-48_ or *bla*_OXA-244_ were located on different genetic elements with variable sizes of ∼2,6 kb, ∼11 kb or ∼20 kb. The *bla*_OXA-48_-like genetic element was flanked by IS1 family transposase IS1R and IS1D and had the IS1R-IS1D −*bla*_OXA-48_-insert-IS1R-IS1D structure or variants thereof (Fig. 5b). The sizes of the genetic elements was determined as the sequence in between the flanking IS1R and IS1D, thereby excluding the size of the IS1R/1D sequence. The chromosomal insertion sites of *bla*_OXA-48_-like genes and length of the insertion element varied per sequence type. The *bla*_OXA-244_ allele was not found in the ST127 genetic background. In *K. pneumoniae bla*_OXA-48_ was also found to be embedded between two IS4 family transposase IS10A genes. Comparison of pOXA-48 plasmids with the chromosomal *bla*_OXA-48_ insertions, revealed that these chromosomal insertions resembled variable regions of plasmid region 1 (Fig. 5b). A 15-nucleotide direct repeat (DR) GGTAATGACTCCAAC was typically located directly upstream IS1R thereby flanking the *bla*_OXA-48_-like insertion element. This DR sequence occurs on average 1 to 2x in pOXA-48-1 to pOXA-48-5 plasmids, except in pOXA-181 and pOXA-232 plasmids (Fig. 5c). The DR was found on average 9x in the 47 *E. coli* chromosomes with *bla*_OXA-48_-like, compared to 4.6x the 5 *K. pneumoniae* chromosomes containing *bla*_OXA-48_. The DR occurred on average 9x, 8x, and 4x in *E. coli* ST38, ST69 and ST127, respectively. In only four of the 52 chromosomes analyzed, the *bla*_OXA-48_ region was flanked by one single DR sequence if the orientation of the carbapenemase allele was in reverse orientation (Table 3). In one chromosome, no DR sequence or truncates hereof were found. In *K. pneumoniae*, in two ST45 of the four isolates with *bla*_OXA-48_ was inserted in the same location in the chromosome through a highly comparable genetic element (Fig. 5b). In a more distantly related *K. pneumoniae* ST101 isolate, a mobile genetic element of ∼2.4 kb *bla*_OXA-48_ was localized in a distinct region, as also for the chromosomes retrieved from NCBI (Fig. 5b).

**Table 3.**
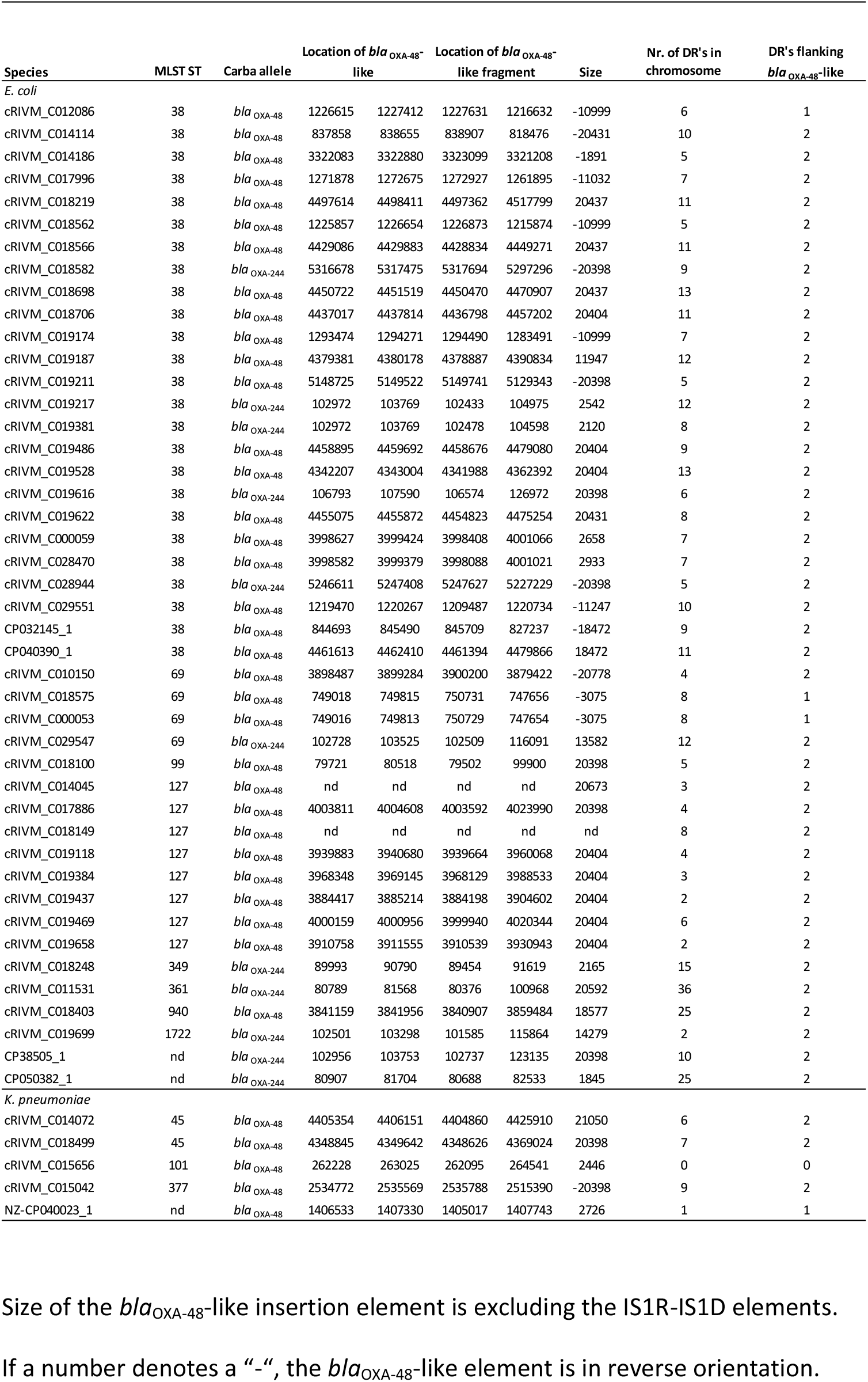
Characteristics of the *bla*_OXA-48_-like chromosomal insertion site, direct repeat and insertion element.

| Species | MLST ST | Carba allele | Location of *bla*<sub>OXA-48</sub>-like |  | Location of *bla*<sub>OXA-48</sub>-like fragment |  | Size | Nr. of DR's in chromosome | DR's flanking *bla*<sub>OXA-48</sub>-like |
| --- | --- | --- | --- | --- | --- | --- | --- | --- | --- |
| *E. coli* |  |  |  |  |  |  |  |  |  |
| cRIVM_C012086 | 38 | *bla*<sub>OXA-48</sub> | 1226615 | 1227412 | 1227631 | 1216632 | -10999 | 6 | 1 |
| cRIVM_C014114 | 38 | *bla*<sub>OXA-48</sub> | 837858 | 838655 | 838907 | 818476 | -20431 | 10 | 2 |
| cRIVM_C014186 | 38 | *bla*<sub>OXA-48</sub> | 3322083 | 3322880 | 3323099 | 3321208 | -1891 | 5 | 2 |
| cRIVM_C017996 | 38 | *bla*<sub>OXA-48</sub> | 1271878 | 1272675 | 1272927 | 1261895 | -11032 | 7 | 2 |
| cRIVM_C018219 | 38 | *bla*<sub>OXA-48</sub> | 4497614 | 4498411 | 4497362 | 4517799 | 20437 | 11 | 2 |
| cRIVM_C018562 | 38 | *bla*<sub>OXA-48</sub> | 1225857 | 1226654 | 1226873 | 1215874 | -10999 | 5 | 2 |
| cRIVM_C018566 | 38 | *bla*<sub>OXA-48</sub> | 4429086 | 4429883 | 4428834 | 4449271 | 20437 | 11 | 2 |
| cRIVM_C018582 | 38 | *bla*<sub>OXA-244</sub> | 5316678 | 5317475 | 5317694 | 5297296 | -20398 | 9 | 2 |
| cRIVM_C018698 | 38 | *bla*<sub>OXA-48</sub> | 4450722 | 4451519 | 4450470 | 4470907 | 20437 | 13 | 2 |
| cRIVM_C018706 | 38 | *bla*<sub>OXA-48</sub> | 4437017 | 4437814 | 4436798 | 4457202 | 20404 | 11 | 2 |
| cRIVM_C019174 | 38 | *bla*<sub>OXA-48</sub> | 1293474 | 1294271 | 1294490 | 1283491 | -10999 | 7 | 2 |
| cRIVM_C019187 | 38 | *bla*<sub>OXA-48</sub> | 4379381 | 4380178 | 4378887 | 4390834 | 11947 | 12 | 2 |
| cRIVM_C019211 | 38 | *bla*<sub>OXA-48</sub> | 5148725 | 5149522 | 5149741 | 5129343 | -20398 | 5 | 2 |
| cRIVM_C019217 | 38 | *bla*<sub>OXA-244</sub> | 102972 | 103769 | 102433 | 104975 | 2542 | 12 | 2 |
| cRIVM_C019381 | 38 | *bla*<sub>OXA-244</sub> | 102972 | 103769 | 102478 | 104598 | 2120 | 8 | 2 |
| cRIVM_C019486 | 38 | *bla*<sub>OXA-48</sub> | 4458895 | 4459692 | 4458676 | 4479080 | 20404 | 9 | 2 |
| cRIVM_C019528 | 38 | *bla*<sub>OXA-48</sub> | 4342207 | 4343004 | 4341988 | 4362392 | 20404 | 13 | 2 |
| cRIVM_C019616 | 38 | *bla*<sub>OXA-244</sub> | 106793 | 107590 | 106574 | 126972 | 20398 | 6 | 2 |
| cRIVM_C019622 | 38 | *bla*<sub>OXA-48</sub> | 4455075 | 4455872 | 4454823 | 4475254 | 20431 | 8 | 2 |
| cRIVM_C000059 | 38 | *bla*<sub>OXA-48</sub> | 3998627 | 3999424 | 3998408 | 4001066 | 2658 | 7 | 2 |
| cRIVM_C028470 | 38 | *bla*<sub>OXA-48</sub> | 3998582 | 3999379 | 3998088 | 4001021 | 2933 | 7 | 2 |
| cRIVM_C028944 | 38 | *bla*<sub>OXA-244</sub> | 5246611 | 5247408 | 5247627 | 5227229 | -20398 | 5 | 2 |
| cRIVM_C029551 | 38 | *bla*<sub>OXA-48</sub> | 1219470 | 1220267 | 1209487 | 1220734 | -11247 | 10 | 2 |
| CP032145_1 | 38 | *bla*<sub>OXA-48</sub> | 844693 | 845490 | 845709 | 827237 | -18472 | 9 | 2 |
| CP040390_1 | 38 | *bla*<sub>OXA-48</sub> | 4461613 | 4462410 | 4461394 | 4479866 | 18472 | 11 | 2 |
| cRIVM_C010150 | 69 | *bla*<sub>OXA-48</sub> | 3898487 | 3899284 | 3900200 | 3879422 | -20778 | 4 | 2 |
| cRIVM_C018575 | 69 | *bla*<sub>OXA-48</sub> | 749018 | 749815 | 750731 | 747656 | -3075 | 8 | 1 |
| cRIVM_C000053 | 69 | *bla*<sub>OXA-48</sub> | 749016 | 749813 | 750729 | 747654 | -3075 | 8 | 1 |
| cRIVM_C029547 | 69 | *bla*<sub>OXA-244</sub> | 102728 | 103525 | 102509 | 116091 | 13582 | 12 | 2 |
| cRIVM_C018100 | 99 | *bla*<sub>OXA-48</sub> | 79721 | 80518 | 79502 | 99900 | 20398 | 5 | 2 |
| cRIVM_C014045 | 127 | *bla*<sub>OXA-48</sub> | nd | nd | nd | nd | 20673 | 3 | 2 |
| cRIVM_C017886 | 127 | *bla*<sub>OXA-48</sub> | 4003811 | 4004608 | 4003592 | 4023990 | 20398 | 4 | 2 |
| cRIVM_C018149 | 127 | *bla*<sub>OXA-48</sub> | nd | nd | nd | nd | nd | 8 | 2 |
| cRIVM_C019118 | 127 | *bla*<sub>OXA-48</sub> | 3939883 | 3940680 | 3939664 | 3960068 | 20404 | 4 | 2 |
| cRIVM_C019384 | 127 | *bla*<sub>OXA-48</sub> | 3968348 | 3969145 | 3968129 | 3988533 | 20404 | 3 | 2 |
| cRIVM_C019437 | 127 | *bla*<sub>OXA-48</sub> | 3884417 | 3885214 | 3884198 | 3904602 | 20404 | 2 | 2 |
| cRIVM_C019469 | 127 | *bla*<sub>OXA-48</sub> | 4000159 | 4000956 | 3999940 | 4020344 | 20404 | 6 | 2 |
| cRIVM_C019658 | 127 | *bla*<sub>OXA-48</sub> | 3910758 | 3911555 | 3910539 | 3930943 | 20404 | 2 | 2 |
| cRIVM_C018248 | 349 | *bla*<sub>OXA-244</sub> | 89993 | 90790 | 89454 | 91619 | 2165 | 15 | 2 |
| cRIVM_C011531 | 361 | *bla*<sub>OXA-244</sub> | 80789 | 81568 | 80376 | 100968 | 20592 | 36 | 2 |
| cRIVM_C018403 | 940 | *bla*<sub>OXA-48</sub> | 3841159 | 3841956 | 3840907 | 3859484 | 18577 | 25 | 2 |
| cRIVM_C019699 | 1722 | *bla*<sub>OXA-244</sub> | 102501 | 103298 | 101585 | 115864 | 14279 | 2 | 2 |
| CP38505_1 | nd | *bla*<sub>OXA-244</sub> | 102956 | 103753 | 102737 | 123135 | 20398 | 10 | 2 |
| CP050382_1 | nd | *bla*<sub>OXA-244</sub> | 80907 | 81704 | 80688 | 82533 | 1845 | 25 | 2 |
| *K. pneumoniae* |  |  |  |  |  |  |  |  |  |
| cRIVM_C014072 | 45 | *bla*<sub>OXA-48</sub> | 4405354 | 4406151 | 4404860 | 4425910 | 21050 | 6 | 2 |
| cRIVM_C018499 | 45 | *bla*<sub>OXA-48</sub> | 4348845 | 4349642 | 4348626 | 4369024 | 20398 | 7 | 2 |
| cRIVM_C015656 | 101 | *bla*<sub>OXA-48</sub> | 262228 | 263025 | 262095 | 264541 | 2446 | 0 | 0 |
| cRIVM_C015042 | 377 | *bla*<sub>OXA-48</sub> | 2534772 | 2535569 | 2535788 | 2515390 | -20398 | 9 | 2 |
| NZ-CP040023_1 | nd | *bla*<sub>OXA-48</sub> | 1406533 | 1407330 | 1405017 | 1407743 | 2726 | 1 | 1 |

Size of the *bla*<sub>OXA-48</sub>-like insertion element is excluding the IS1R-IS1D elements.
If a number denotes a “-”, the *bla*<sub>OXA-48</sub>-like element is in reverse orientation.

**Figure 5.**
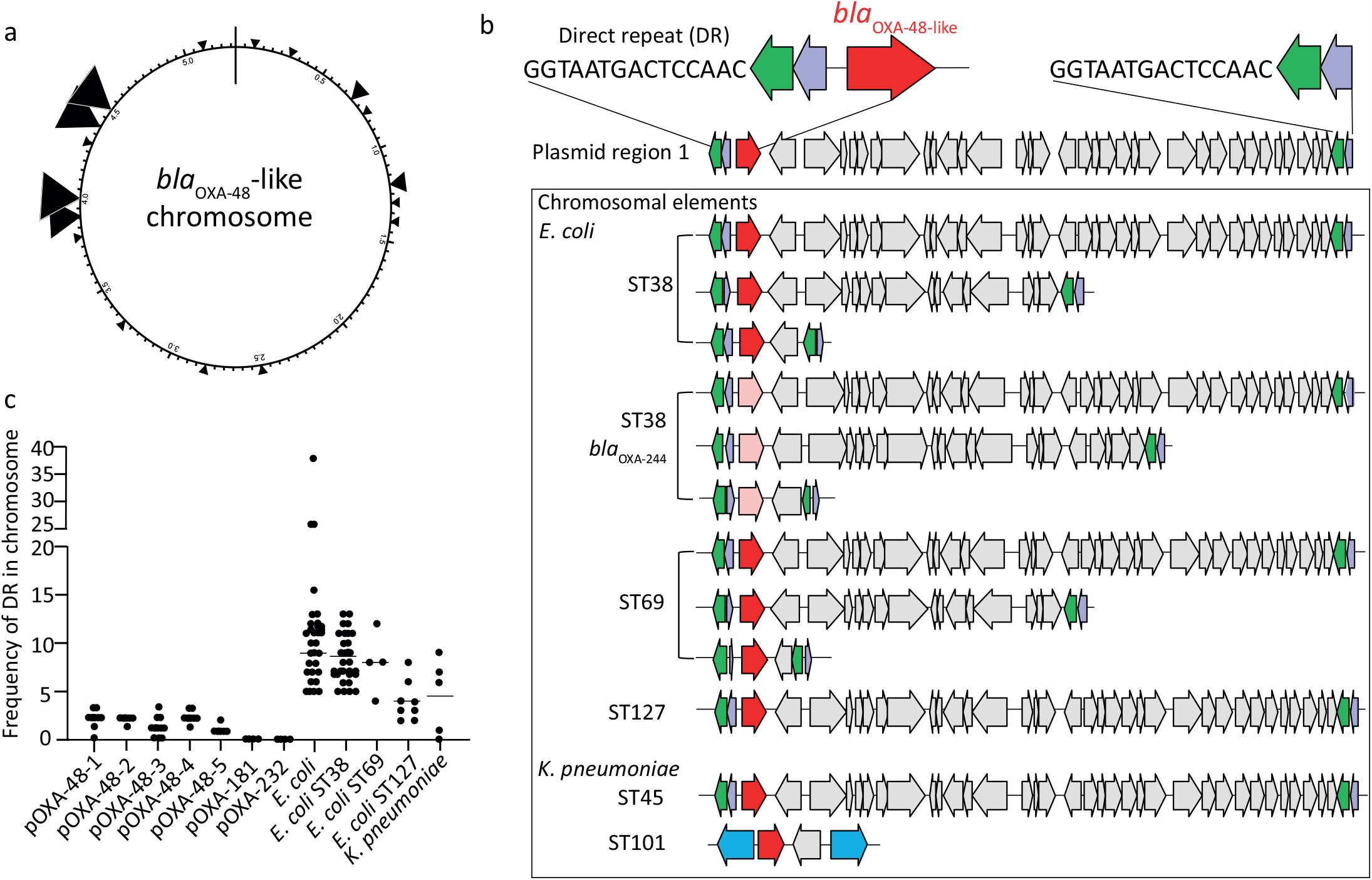
Distinct integration sites of variable *bla*_OXA-48_ and *bla*_OXA-244_ elements in the chromosome. (a) Artificial chromosome in which the different *bla*_OXA-48_-like insertion positions are indicated by triangles.(b) Comparison of plasmid region 1 with chromosomal insertion sites of *bla*_OXA-48_-like. Arrows indicate open reading frames of which *bla*_OXA-48_ is depicted in red and *bla*_OXA-244_ in light red. DR indicated the direct repeat sequence GGTAATGACTCCAAC located upstream IS1R. Sequence types are depicted by ST and sizes of the different insertion sequences are indicated in kilobase (kb). (c) Frequency of the DR sequence in *bla*_OXA-48_-like plasmids and chromosomes of *E. coli* and *K. pneumoniae*.

## Discussion

We dissected the architecture of 179 complete plasmids carrying *bla*_OXA-48_-like alleles and 44 *bla*_OXA-48_-like containing chromosomes of *E. coli* and *K. pneumoniae* isolates obtained from the Dutch national CPE surveillance program in comparison with *bla*_OXA-48_-like plasmids and chromosomes reported in the NCBI databank. The overall *bla*_OXA-48_-like plasmid population in the Netherlands is conserved and compares to internationally reported plasmids. Most of the *bla*_OXA-48_-like plasmids from both *E. coli* and *K. pneumoniae* could be clustered into seven distinct genotypic plasmid groups, which were characterized by a paucity in AMR genes, marked differences in gene content, replicon family, size, and %G+C content. This suggests the plasmids studied here have distinct origins and transferred horizontally among CPE world-wide. In contrast to pOXA-181 and pOXA-232 plasmids, which were highly conserved, a group of pOXA-48 plasmids were diverse in genetic composition with sequence variation as high as 20%. The presence of a variety of transposases and IS sequences, in addition to conjugation machinery may attributed to the genetic diversity of the pOXA-48 plasmids, in particular in the pOXA-48-3 and pOXA-48-5 plasmid subgroups.

There was an additional group of genetically highly diverse *bla*_OXA-48_-like plasmids obtained in the Netherlands with a large range in %G+C content, a variety of IncL and non-IncL-type replicons (IncR, IncFII or IncY), AMR genes, and low inter-plasmid similarity. This suggests the presence of a potentially recently introduced set of plasmids that have not yet been widely spread in the Netherlands. OXA-48 plasmids with either an IncR, IncFII or IncY replicon have only recently been described and are relatively rare (20–22). *bla*_OXA-48_-like plasmids occurred in globally disseminated *E. coli* and *K. pneumoniae* isolates with known genetic backgrounds such as *E. coli* ST38, and *K. pneumoniae* ST307, but also multiple new STs demonstrating continuous dissemination of antimicrobial resistance plasmids to new genetic backgrounds. To date, no double combinations of *bla*_OXA-48_-like of alleles have been detected in one strain although combinations with other carbapenemase alleles such as either *bla*_NDM-1_ or *bla*_NDM-5_ exist.

In this study we also detected chromosomally localized *bla*_OXA-48_ and *bla*_OXA-244_ alleles, but not chromosomal *bla*_OXA-181_ and *bla*_OXA-232_ alleles. This is in contrast to reports from other countries, where chromosomally localized *bla*_OXA-181_ and *bla*_OXA-232_ alleles have been found occasionally (6, 23). Chromosomal insertion of *bla*_OXA-48_ or *bla*_OXA-244_ may have occurred through IS1R-mediated transposition and recombination of OXA-48 plasmid sequences into *E. coli* and *K. pneumoniae* chromosomes with distinct genetic compositions (15). The various lengths and compositions of *bla*_OXA-48_-like segments and a variety of locations in the chromosome, suggests that multiple transposition and recombination events have occurred. The chromosomal *bla*_OXA-48_ segment likely originated from plasmids belonging to the pOXA-48-1 to pOXA-48-5 groups. A potential insertion target site, a 15-bp direct repeat, was present in multiple copies in the chromosome and was found only in pOXA-48-1 to pOXA-48-5 plasmids, but not in pOXA-181 and pOXA-232 plasmids. This direct repeat was also found more frequently in *E. coli* than in *K. pneumoniae* chromosomes, which may explain why more *E. coli* than *K. pneumoniae* isolates harbor chromosomal *bla*_OXA-48_/*bla*_OXA-244_ and not *bla*_OXA-181_/*bla*_OXA-232_.

The majority of the *bla*_OXA-48_ containing *K. pneumoniae* isolates in this study had MICs for meropenem above the clinical breakpoint in contrast to *E. coli*, which were mostly sensitive. The distinct *bla*_OXA-48_-like alleles had different meropenem susceptibilities in *K. pneumoniae* and *E. coli* isolates, indicating that not each allele results in the same resistance phenotype. Especially *K. pneumoniae* containing *bla*_OXA-232_ were highly resistant, which can possibly be attributed to a high copy number of pOXA-232 plasmids (24). Alternatively, OXA-48 enzyme production, an altered affinity for meropenem, or other determinants such as porins, efflux pumps or the presence of additional extended-spectrum β-lactamases can be responsible for this phenomenon as well (25, 26).

In conclusion, long-read sequencing of isolates from the Dutch National CPE surveillance contributed to the dissection of the architecture of *bla*_OXA-48_-like plasmids and *bla*_OXA-48_-like chromosome insertions of CPE in the Netherlands. Conjugation machinery, transposable elements, and/or virulence determinants may contribute to plasmid diversification and dissemination, and represent important features that warrant future investigation. Additional long-read sequencing efforts of plasmids of CPE are required to monitor the changing plasmid reservoir involved in the spread of antibiotic resistance determinants in the Netherlands and beyond.

## Materials and methods

### Bacterial isolates

For the Dutch National CPE Surveillance program, medical microbiology laboratories from the Netherlands routinely send *Enterobacterales* isolates with a meropenem minimum inhibitory concentration (MIC) of >0.25 mg/L and/or an imipenem MIC of >1 mg/L or genotypical or phenotypical evidence of carbapenemase production to the National Institute of Public Health and the Environment through Type-Ned, an online platform (3). The low MIC threshold for submission was chosen to monitor CPE instead of carbapenem-resistant *Enterobacterales* (CRE), because CPE represent a reservoir for the spread of antibiotic resistance genes. In this study, 537 carbapenemase-producing *E. coli* and *K. pneumoniae* isolates carrying *bla*_OXA-48_-like alleles (*bla*_OXA-48_, *bla*_OXA-181_, *bla*_OXA-232_) were included and were collected from January 1^st^ 2014 until December 31^st^ 2019 (Table S1, Suppl. File 1). Only the first submitted *E. coli* or *K. pneumoniae* isolate with a *bla*_OXA-48_-like allele per person in this study period was included.

### Antimicrobial susceptibility testing

Resistance to carbapenem was confirmed by assessing the MIC for meropenem using an Etest (bioMérieux Inc., Marcy l’Etoile, France). Based on the clinical breakpoints according to EUCAST, the isolates were classified as sensitive (≤2 mg/L), intermediate (>2 mg/L and ≤8 mg/L) and resistant (>8 mg/L) to meropenem. Isolates were analyzed for carbapenemase production using the carbapenem inactivation method (CIM) (27).

### Next-generation sequencing

*E. coli* and *K. pneumoniae* isolates were subjected to next-generation sequencing (NGS) using the Illumina HiSeq 2500 (BaseClear, Leiden, the Netherlands). The antibiotic resistance gene profile and plasmid replicon compositions in all of the isolates were determined by interrogating the ResFinder (version 3.1.0) and PlasmidFinder (version 2.0.2) databases available from the Center for Genomic Epidemiology (28, 29). For ResFinder, a 90% identity threshold and a minimum length of 60% were used as criteria, whereas for PlasmidFinder, an identity of 95% was utilized. The resulting NGS derived data, such as resistance genes, replicons, wgMLST profiles were imported into BioNumerics version 7.6.3 for subsequent comparative analyses (Applied Maths, Sint-Martens-Latem, Belgium).

### Long-read third-generation sequencing

High molecular weight DNA was isolated using an in-house developed protocol as described previously (3). The Oxford Nanopore protocol SQK-LSK108 (https://community.nanoporetech.com) and the expansion kit for native barcoding EXP-NBD104 was used. A shearing step was performed using g-TUBE’s (Covaris) to obtain an average DNA fragment size of 8 kb for isolates from 2014-2018. This shearing step was omitted for isolates from 2019. The DNA was repaired using FFPE and end-repair kits (New England BioLabs) followed by ligation of barcodes with bead clean up using AMPure XP (Beckman Coulter) after each step. Barcoded isolates were pooled and sequencing adapters were added by ligation. The final library was loaded onto a MinION flow cell (MIN-106 R9.4.1). The 48-hour sequence run was started without live base calling enabled on a MinION device connected to a desktop computer. After the sequence run, base calling and de-multiplexing was performed using Albacore 2.3.1 and a single FASTA file per isolate was extracted from the FAST5 files using Poretools 0.5.1 (30). Illumina and Nanopore data were used in a hybrid assembly performed by Unicycler v0.4.4 (31). The resulting contig files were annotated using Prokka and were subsequently loaded into BioNumerics for further analyses (32).

### Plasmid content analysis

For annotation a Conda environment was set up with packages to facilitate a Snakemake pipeline which could process samples in bulk, preform initial annotation with Prokka and enhancement with BLAST+ (33, 34). Prokka annotation was executed in two stages, in the first stage it identified the coordinates of candidate genes with Prodigal, in the second step it predicted these genes by utilizing user set databases and its default the Swissprot database (35, 36). To preserve the speed of the initial annotation we prepared a small database by combining sequence data from the ResFinder database and the PlasmidFinder database (28, 29). If Prokka was unable to predict a gene it will label the coordinate as a hypothetical protein. In order to reduce the hypothetical proteins in our annotation we used a set of custom Python scripts to extract and prepare them for BLAST+. After alignment with BLAST+ the supplemented Python code was used to replace the hypothetical proteins in the initial annotation file with their best alignment match (Suppl. File 2). BioNumerics was used to extract and analyze the presence of annotated genes and tranposases in the different plasmids. The data was plotted, analyzed and visualized in Excel.

### Plasmid and chromosome comparisons

BioNumerics was used to compare complete plasmid DNA sequence and circular and linear chromosome datasets. Linear assembly contigs were omitted in this study. The CLC Genomics Workbench version 12.0 software (www.qiagenbioinformatics.com) was used to retrieve *bla*_OXA-48_-like plasmids and chromosomes from NCBI (Table S1). These plasmids and chromosomes were stripped from their annotations and re-annotated again using Prokka. All chromosomes have the *dnaA* gene as starting point in order to determine relative locations of *bla*_OXA-48_-like alleles. For analysis of the plasmid gene content, the *bla*_OXA-48_ or *bla*_OXA-48_-like allele was set as the starting point.

### Ethics statement

The bacterial isolates belong to the medical microbiological laboratories participating in the Dutch National CPE Surveillance program and were obtained as part of routine clinical care in the past years. Since no identifiable personal data were collected and data were analyzed and processed anonymously, written or verbal patient consent was not required. According to the Dutch Medical Research Involving Human Subjects Act (WMO) this study was exempt from review by an Institutional Review Board.

### Minimum Spanning Tree and UPGMA analyses

The BioNumerics software was used to generate a minimum spanning tree (MST) or an UPGMA hierarchical clustering as described previously (3). The MST was based on in-house *E. coli* and *K. pneumoniae* wgMLST schemes. The categorical coefficient was used to calculate the MST.

### NGS and plasmid data availability

The Illumina (NGS) sequence data set generated and analyzed in this study are available in NCBI in the European Nucleotide Archive (ENA) under project number xxx. The plasmid sequences are deposited in Genbank and available through the accession numbers xxx.

## Acknowledgements

We thank all the members of the Dutch CPE surveillance study Group and the Dutch medical microbiology laboratories for submitting CPE isolates to the RIVM for the national CPE surveillance program. We also thank Prof. Dr. E. Kuijper, Dr. S.C. de Greeff and Dr. D.W. Notermans for critical reading of this manuscript. Members of the Dutch CPE surveillance Study Group:

A. Maijer-Reuwer, ADRZ medisch centrum, Department of Medical Microbiology, Goes

M.A. Leversteijn-van Hall, Alrijne Hospital, Department of Medical Microbiology, Leiden

J.A.J.W. Kluytmans, Amphia Hospital, Microvida Laboratory for Microbiology, Breda

I.J.B. Spijkerman, Amsterdam UMC - location AMC, Department of Medical Microbiology, Amsterdam

K. van Dijk, Amsterdam UMC - location Vumc, Department of Medical Microbiology and Infection Control, Amsterdam

T. Halaby, Analytical Diagnostic Center N.V. Curaçao, Department of Medical Microbiology, Willemstad (Curaçao)

B. Zwart, Atalmedial, Department of Medical Microbiology, Amsterdam

B.M.W. Diederen, Bravis Hospital/ZorgSaam Hospital Zeeuws-Vlaanderen, Department of Medical Microbiology, Roosendaal/Terneuzen

A. Voss, Canisius Wilhelmina Hospital, Department of Medical Microbiology and Infectious Diseases, Nijmegen

J.W. Dorigo-Zetsma, CBSL, Department of Medical Microbiology, Hilversum

D.W. Notermans, Centre for Infectious Disease Control, National Institute for Public Health and the Environment, Bilthoven

A. Ott, Certe, Department of Medical Microbiology, Groningen

J.H. Oudbier, Comicro, Department of Medical Microbiology, Hoorn

M. van der Vusse, Deventer Hospital, Department of Medical Microbiology, Deventer

A.L.M. Vlek, Diakonessenhuis, Department of Medical Microbiology and Immunology, Utrecht

A.G.M. Buiting, Elisabeth-TweeSteden (ETZ) Hospital, Department of Medical Microbiology and Immunology, Tilburg

L. Bode, Erasmus University Medical Center, Department of Medical Microbiology, Rotterdam

S. Paltansing, Franciscus Gasthuis & Vlietland, Department of Medical Microbiology and Infection Control, Rotterdam

A.J. van Griethuysen, Gelderse Vallei Hospital, Department of Medical Microbiology, Ede

M. den Reijer, Gelre Hospitals, Department of Medical Microbiology and Infection prevention, Apeldoorn

M. van Trijp, Groene Hart Hospital, Department of Medical Microbiology and Infection Prevention, Gouda

E.P.M. van Elzakker, Haga Hospital, Department of Medical Microbiology, ‘s-Gravenhage

A.E. Muller, HMC Westeinde Hospital, Department of Medical Microbiology, ‘s-Gravenhage

M.P.M. van der Linden, IJsselland hospital, Department of Medical Microbiology, Capelle a/d IJssel

M. van Rijn, Ikazia Hospital, Department of Medical Microbiology, Rotterdam

M.J.H.M. Wolfhagen, Isala Hospital, Laboratory of Medical Microbiology and Infectious Diseases, Zwolle

K. Waar, Izore Centre for Infectious Diseases Friesland, Department of Medical Microbiology, Leeuwarden

E. Kolwijck, Jeroen Bosch Hospital, Department of Medical Microbiology and Infection Control, ‘s-Hertogenbosch

W. Silvis, LabMicTA, Regional Laboratory of Microbiology Twente Achterhoek, Hengelo

T. Schulin, Laurentius Hospital, Department of Medical Microbiology, Roermond

M. Damen, Maasstad Hospital, Department of Medical Microbiology, Rotterdam

S. Dinant, Maasstad Hospital, Department of Medical Microbiology, Rotterdam

S.P. van Mens, Maastricht University Medical Centre, Department of Medical Microbiology, Maastricht

D.C. Melles, Meander Medical Center, Department of Medical Microbiology, Amersfoort

J.W.T. Cohen Stuart, Noordwest Ziekenhuisgroep, Department of Medical Microbiology, Alkmaar

M.L. van Ogtrop, Onze Lieve Vrouwe Gasthuis, Department of Medical Microbiology, Amsterdam

I.T.M.A. Overdevest, PAMM, Department of Medical Microbiology, Veldhoven

A.P. van Dam, Amsterdam Health Service, Public Health Laboratory, Amsterdam

H. Wertheim, Radboud University Medical Center, Department of Medical Microbiology, Nijmegen

H.M.E. Frénay, Regional Laboratory Medical Microbiology (RLM), Department of Medical Microbiology, Dordrecht

J.C. Sinnige, Regional Laboratory of Public Health, Department of Medical Microbiology, Haarlem

E.E. Mattsson, Reinier de Graaf Groep, Department of Medical Microbiology, Delft

R.W. Bosboom, Rijnstate Hospital, Laboratory for Medical Microbiology and Immunology, Velp

A. Stam, Saltro Diagnostic Centre, Department of Medical Microbiology, Utrecht

E. de Jong, Slingeland Hospital, Department of Medical Microbiology, Doetinchem

N. Roescher, St Antonius Hospital, Department of Medical Microbiology and Immunology, Nieuwegein

E. Heikens, St Jansdal Hospital, Department of Medical Microbiology, Harderwijk

R. Steingrover, St. Maarten Laboratory Services, Department of Medical Microbiology, Cay Hill (St. Maarten)

A. Troelstra, University Medical Center Utrecht, Department of Medical Microbiology, Utrecht

E. Bathoorn, University of Groningen, Department of Medical Microbiology, Groningen

T.A.M. Trienekens, VieCuri Medical Center, Department of Medical Microbiology, Venlo

D.W. van Dam, Zuyderland Medical Centre, Department of Medical Microbiology and Infection Control, Sittard-Geleen

E.I.G.B. de Brauwer, Zuyderland Medical Centre, Department of Medical Microbiology and Infection Control, Heerlen

F.S. Stals, Zuyderland Medical Centre, Department of Medical Microbiology and Infection Control, Heerlen

---

**Table S1.** CPE analyzed in this study.

| Year | *E. coli* | *K. pneumoniae* | Total |
| --- | --- | --- | --- |
| 2014 | 7 | 13 | 20 |
| 2015 | 15 | 36 | 51 |
| 2016 | 30 | 43 | 73 |
| 2017 | 38 | 51 | 89 |
| 2018 | 58 | 87 | 145 |
| 2019 | 82 | 77 | 159 |
| Total | 230 | 307 | 537 |

**Table S2.**
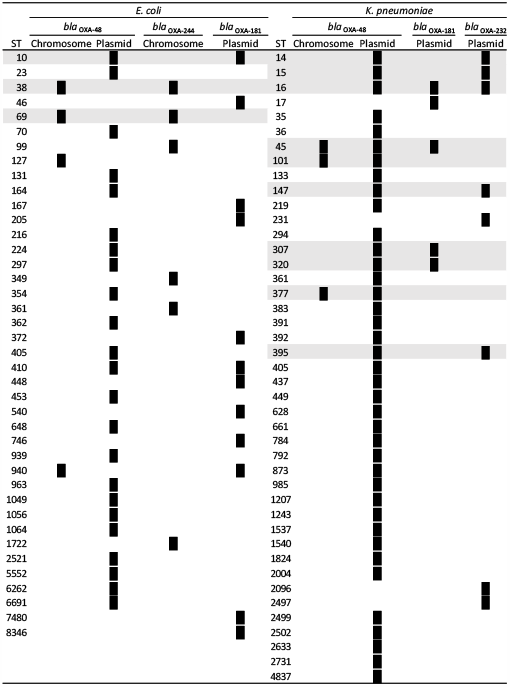
Classical MLST sequence types of the isolates analyzed in this study.

